# Machine learning models for predicting protein condensate formation from sequence determinants and embeddings

**DOI:** 10.1101/2020.10.26.354753

**Authors:** Kadi L. Saar, Alexey S. Morgunov, Runzhang Qi, William E. Arter, Georg Krainer, Alpha A. Lee, Tuomas P. J. Knowles

## Abstract

Intracellular phase separation of proteins into biomolecular condensates is increasingly recognised as an important phenomenon for cellular compartmentalisation and regulation of biological function. Different hypotheses about the parameters that determine the tendency of proteins to form condensates have been proposed with some of them probed experimentally through the use of constructs generated by sequence alterations. To broaden the scope of these observations, here, we established an *in silico* strategy for understanding on a global level the associations between protein sequence and condensate formation, and used this information to construct machine learning classifiers for predicting liquid–liquid phase separation (LLPS) from protein sequence. Our analysis highlighted that LLPS–prone sequences are more disordered, hydrophobic and of lower Shannon entropy than sequences in the Protein Data Bank or the Swiss-Prot database, and have their disordered regions enriched in polar, aromatic and charged residues. Using these determining features together with neural network based word2vec sequence embeddings, we developed machine learning classifiers for predicting protein condensate formation. Our model, trained to distinguish LLPS-prone sequences from structured proteins, achieved high accuracy (93%; 25-fold cross-validation) and identified condensate forming sequences from external independent test data at 97% sensitivity. Moreover, in combination with a classifier that had developed a nuanced insight into the features governing protein phase behaviour by learning to distinguish between sequences of varying LLPS propensity, the sensitivity was supplemented with high specificity (approximated ROC–AUC of 0.85). These results provide a platform rooted in molecular principles for understanding protein phase behaviour. The predictor is accessible from https://deephase.ch.cam.ac.uk/.

**Significance Statement:** The tendency of many cellular proteins to form protein-rich biomolecular condensates underlies the formation of subcellular compartments and has been linked to various physiological functions. Understanding the molecular basis of this fundamental process and predicting protein phase behaviour have therefore become important objectives. To develop a global understanding of how protein sequence determines its phase behaviour, here, we constructed bespoke datasets of proteins of varying phase separation propensity and identified explicit biophysical and sequence-specific features common to phase separating proteins. Moreover, by combining this insight with neural network based sequence embeddings, we trained machine learning classifiers that identified phase separating sequences with high accuracy, including from independent external test data. The predictor is available from https://deephase.ch.cam.ac.uk/.

## Introduction

Liquid–liquid phase separation (LLPS) is a widely occurring biomolecular process that can lead to the formation of membraneless organelles within living cells.^1–4^ This process and the resulting condensate bodies are increasingly recognised to play an important role in a wide range of biological processes, including the onset and development of metabolic diseases and neurodegenerative disorders.^5–11^ Understanding the factors that drive the formation of protein-rich biomolecular condensates has thus become an important objective and been the focus of a large number of studies, which have collectively yielded valuable information about the factors that govern protein phase behaviour.^3,4,12,13^

While changes in extrinsic conditions, such as temperature, ionic strength or the level of molecular crowding can strongly modulate LLPS,^14–17^ of fundamental importance to condensate formation is the linear amino acid sequence of a protein, its primary structure. A range of sequence-specific factors governing the formation of protein condensates have been postulated with *π-π* contacts and the valency and patterning of prion-like domains in particular brought forward as central features.^12,13,18–23^ The predictive power of some of these hypothesis has been recently reviewed.^24^ In parallel, studies examining the relationship between protein phase behaviour and its sequence alterations through deletion, truncation and site-specific mutations events have determined various sequence-specific features to be important in modulating the protein phase separation of specific proteins, such as the high abundance of arginine and tyrosine residues in the context of the fused in sarcoma (FUS)-family proteins,^23^ the positioning of tryptophan and other aromatic amino acid residues in TAR DNA-binding protein 43 (TDP-43),^25^ arginine- and glycine-rich disordered domains in LAF–1 protein^26^ and multivalent interactions for the UBQLN2 protein.^27^

To broaden the scope of these observations and understand on a global level the associations between the primary structure of a protein and its tendency to form condensates, here, we developed an *in silico* strategy for analysing the associations between LLPS propensity of a protein and its amino acid sequence and we used this information to construct machine learning classifiers for predicting LLPS propensity from the amino acid sequence. Specifically, by starting with a previously published LLPSDB database collating information on protein phase behaviour under different environmental conditions^28^ and by analysing the concentration under which LLPS had been observed to take place in these experiments, we constructed two datasets including sequences of different LLPS propensity and compared them to fully ordered structures from the Protein Data Bank (PDB)^29^ as well as the Swiss-Prot^30^ database. We observed phase separating proteins to be more hydrophobic, more disordered and of lower Shannon entropy than sequences in these databases. Moreover, by focusing the analysis onto the low complexity regions (LCR), we showed that phase separation propensity correlates with high abundance of polar amino acids within the LCRs, which in turn correlates poorly with that of hydrophobic residues, suggesting that the regions of polar residues are punctuated by only a fixed number of hydrophobic residues regardless of the length of the polar region. These results are in line wit previously proposed models and experimental observations,^18,22^ and add granularity regarding the composition of the LCRs.

Moreover, we combined the outlined sequence-specific characteristics with implicit features based on neural network derived word2vec sequence embeddings and trained classifiers for predicting the propensity of unseen proteins to phase separate. A model trained to distinguish LLPS-prone sequences from structured sequences in the PDB achieved over 93% accuracy (25-fold cross-validation) — a result similar to what has been achieved before.^31^ To further force the model to learn features that could be more informative for predicting LLPS propensity, as opposed to learning to distinguish between highly disjoint sets of proteins showing a stark difference across a variety of features, we trained a second classifier that could discriminate between disordered sequences of different condensate formation propensity. The combination of these strategies enabled us to construct a final model that showed both high specificity and sensitivity (AUC-ROC – 0.85) in identifying LLPS-prone sequences from external data constructed through an exhaustive literature search.^32^ These results shed light onto the physicochemical factors modulating protein condensate formation, and provide a platform rooted in molecular principles for the prediction of protein phase behaviour.

## Results and Discussion

### Construction of datasets and their global sequence comparison

To link the amino acid sequence of a protein to its tendency to form biomolecular condensates, we gathered data from two publicly available datasets, the LLPSDB^28^ and the PDB^29^ and constructed three bespoke datasets — LLPS^+^ and LLPS^-^ comprising sequences with high and low LLPS propensity, respectively, and PDB* consisting of sequences that were very unlikely to phase separate (see below). To create the first two datasets, the LLPSDB dataset was reduced to naturally occurring sequence constructs with no post-translational modifications and further filtered for sequences that were observed to undergo phase separation on their own, in isolation from other components, such as DNA, RNA or an additional protein (see Methods). In the resulting dataset (231 constructs from 120 unique UniProt IDs), the mean concentration at which each construct had been observed to phase separate was estimated and a threshold of 100 μM was used to divide the sequences according to their high (LLPS^+^; 138 constructs; Supplementary Table S1) or low (25 constructs) propensity to phase separate (Figure 1a). The latter sequences were then combined with the remainder of the sequences from the LLPSDB that had not been observed to phase separate homotypically (69 constructs) to form the LLPS^-^ dataset (94 constructs; Supplementary Table S2). We note that this classification does not imply that the proteins in the LLPS^-^ dataset cannot phase separate under any conditions, but it rather allowed us to learn information about sequence-specific features that correlate with enhanced LLPS propensity.

**Figure 1.**
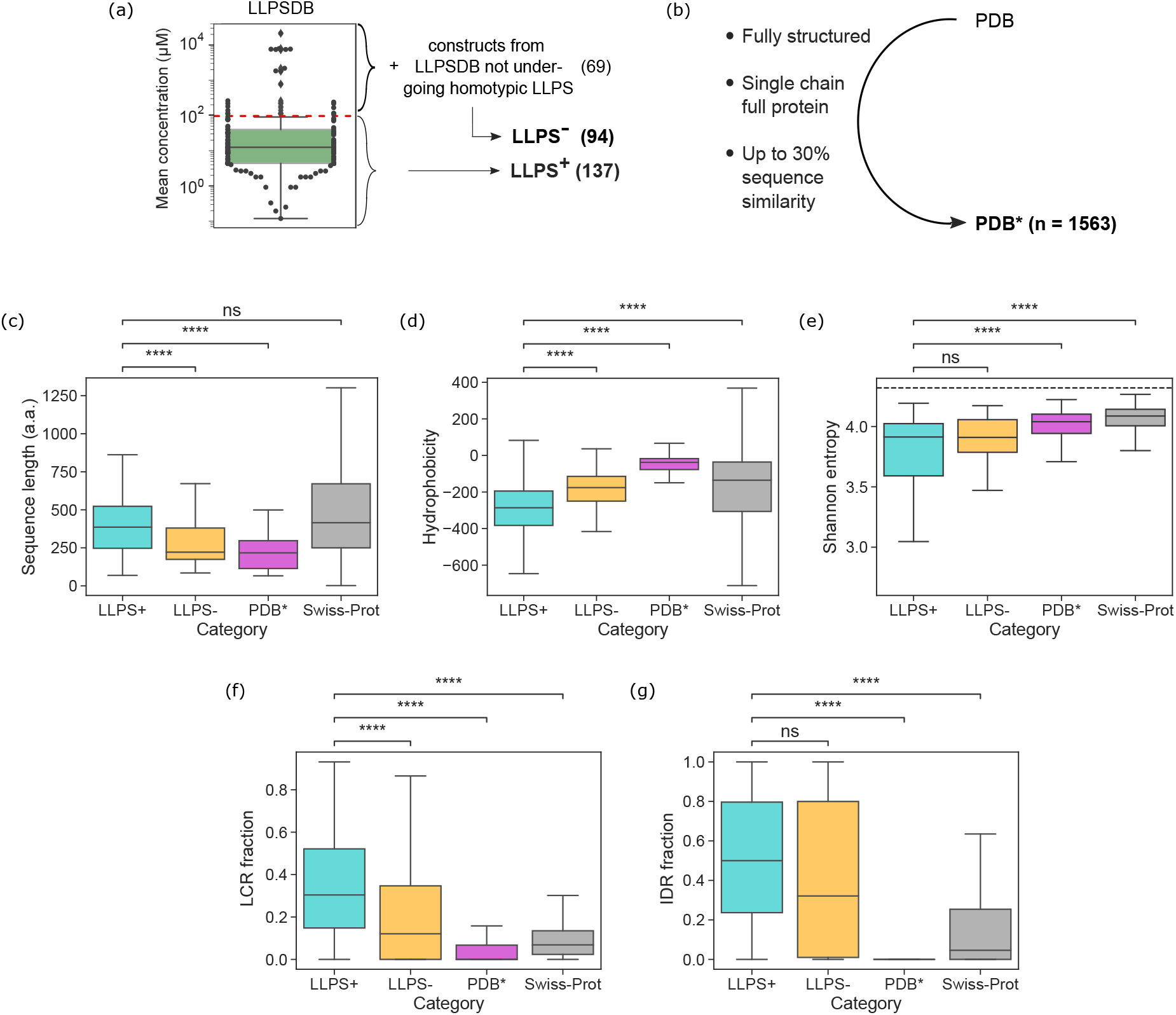
Construction and comparison of the datasets. **(a)** The entries in the LLPSDB dataset^28^ were filtered for single-protein systems and the constructs that phase separated at an average concentration below *c* = 100 μM were classified as having a high propensity to phase-separate (LLPS^+^; 137 constructs) with the remaining 25 constructs together with constructs that had not been observed to phase separate homotypically classified as low propensity dataset (LLPS^-^; 94 constructs). **(b)** The PDB* dataset (1563 constructs) was constructed by filtering the entries in the PDB^29^ to fully structured full-protein single-chains and clustering for sequence similarity (30% cut-off) with a single entry selected from each cluster. **(c)** Sequence length (in amino acids, a.a.), **(d)** hydrophobicity, **(e)** Shannon entropy, the fraction of sequence that is part of **(f)** the low complexity regions (LCR) and **(g)** the intrinsically disordered regions (IDRs) for the three training datasets and the Swiss-Prot. Comparison of the datasets highlighted that LLPS-prone sequences (cyan) were more hydrophobic and of lower Shannon entropy than an average sequence in the Swiss-Prot dataset (grey) and that they had elevated LCR and IDR fractions. The boxes bound data between the upper and the lower quartile, their centre lines indicate the mean value. The ends of the whiskers correspond to values that exceed the boundaries of the interquartile range by 1.5 times its size or to the most extreme value. Significance was tested with Mann-Whitney test, **** denotes a p-value below 10^-4^ and ns denotes no significance at *p* ≤ 0.01. The linear dotted line in panel **(e)** corresponds to the case when all amino acids are present at equal frequencies.

An additional dataset, PDB*, consisting of entries sampled from the PDB was constructed for it to serve as an alternative training set comprising sequences that are highly unlikely to phase separate. This dataset was constructed by including sequences from the PDB that did not include any disordered residues (112,572 chains) filtered for lengths above 50 amino acids with the selected sequences verified via mapping to their UniProt IDs (see Methods). The remaining sequences (13,325) were clustered for sequence identity to retain no more than a single sequence from each cluster (see Methods). This process reduced the original PDB dataset to a diverse set of 1563 fully structured sequences (PDB*; Supplementary Table S3; Figure 1b).

We compared the generated datasets across a range of global sequence-specific features with the aim to understand the factors that are linked with enhanced condensate formation propensity using the Swiss-Prot database^30^ as a reference control (Figure 1c–f). From the analysis, we first concluded that the average construct in the LLPS^+^ dataset (cyan) did not differ in its length from the average construct in Swiss-Prot (grey) but it was longer than the average construct in the LLPS^-^ (orange) or the PDB^*^ dataset (magenta; Figure 1c). We next estimated the hydrophobicity of all the constructs in the four datasets using Kyte and Doolittle hydropathy scale^33^ and concluded LLPS-prone constructs to be less hydrophobic than the sequences in any of the other three datasets (Figure 1d). Finally, we noted that LLPS-prone sequences exhibited lower Shannon entropy than the sequenes in the PDB^*^ or the Swiss-Prot dataset (Figure 1e) — an effect that can be linked to their extended prion-like domains.^22,23^

To understand how sequence complexity and the extent of disorder were linked to the tendency of proteins to undergo phase separation, we employed the SEG algorithm^34^ to extract low complexity regions (LCRs) for all the sequences in the four datasets and the IUPred2 algorithm^35^ to identify their disorded regions (see Methods). This analysis revealed that constructs in the LLPS^+^ dataset had a larger fraction of the sequences that was part of the LCRs than the sequences in any of the other three datasets (Figure 1f) and a higher degree of disorder than sequences in the PDB^*^ or the Swiss-Prot dataset (Figure 1g).

### Amino acid composition of the constructs undergoing LLPS

Having ascertained the length of the low complexity and intrinsically disordered regions as basic parameters that set the constructed datasets apart (Figure 1f-g), we next set out to analyse the amino acid composition of these regions (Figure 2). By classifying the amino acid residues into polar, hydrophobic, aromatic, cationic and anionic categories (see Methods), we observed that the propensity of proteins to undergo LLPS was associated with a higher relative content of polar (blue) and a reduced relative content of hydrophobic (orange) and anionic (purple) residues across the full amino acid sequence (Figure 2a). The increased abundance of polar residues was particularly pronounced within the LCRs with aromatic (green) and cationic (red) residues also being overrepresented within these regions compared to the sequences in the Swiss-Prot database (Figure 2b).

**Figure 2.**
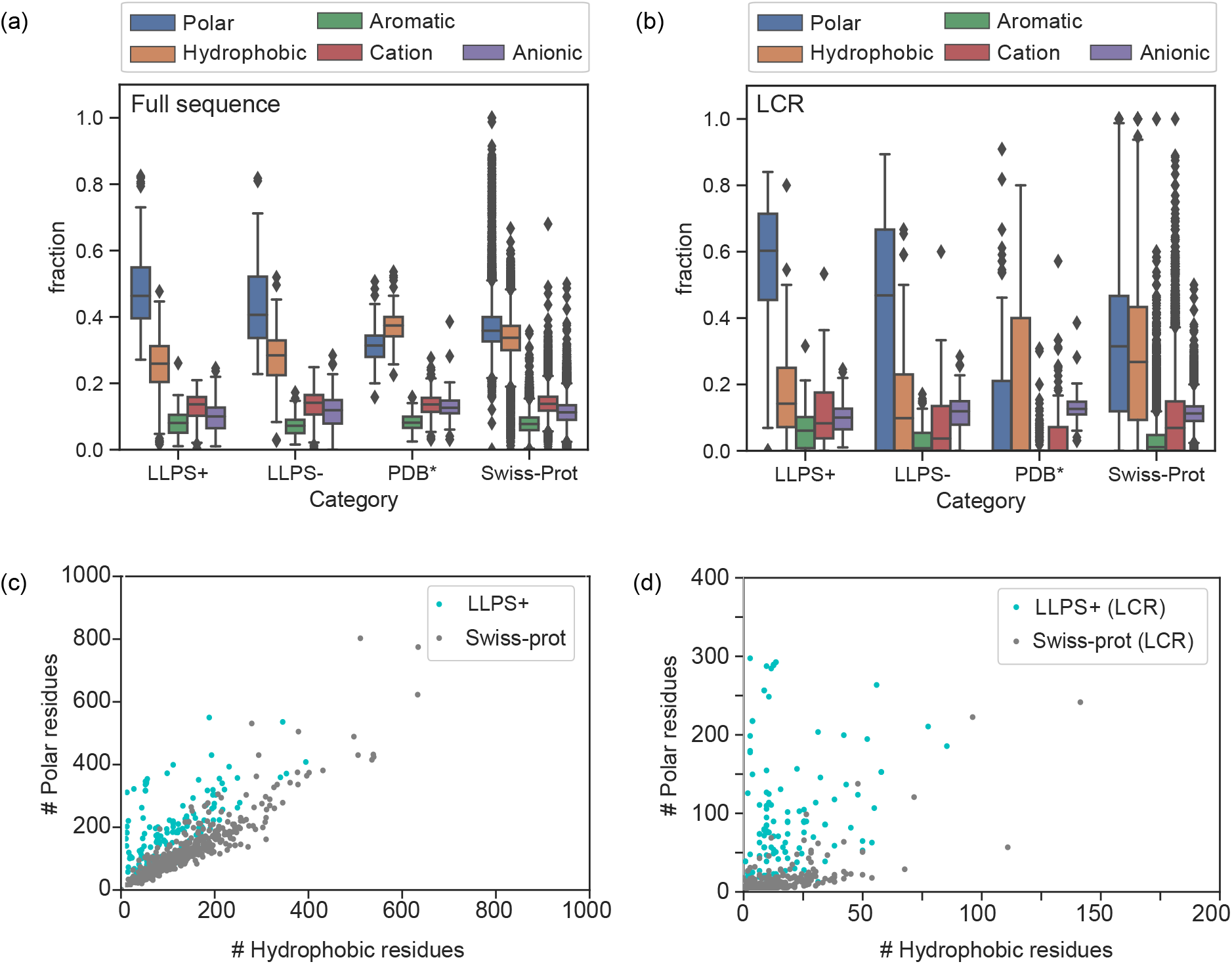
Amino acid compositions of the sequences in the four datasets. **(a)** Comparison of the amino acid composition of the sequences within the LLPS^+^, LLPS^-^, PDB* and the Swiss-Prot dataset highlighted that LLPS^+^ sequences were enriched in polar amino acid residues (blue) and depleted in hydrophobic (orange) and anionic (purple) residues compared to the other three datasets. **(b)** The elevated abundance of polar amino acid residues was particularly prounced within the LCR with the sequences in the LLPS^+^ also having their LCR regions enriched in aromatic (green) and cationic (red) residues compared to the Swiss-Prot database. **(c)** On the level of an individual sequence, polar residues were overrepresented in LLPS-prone sequences with the correlation between the abundance of polar and hydrophobic residues being weaker within the LLPS^+^ dataset than it was in the Swiss-Prot dataset. **(d)** A similar trend was observed within in the LCR. These effects likely originate from the regions of polar residues being punctuated by only a fixed number of hydrophobic residues regardless of the length of the polar region — an observation in line with the stickers-and-spacers model.^22^

We additionally examined the correlation between the abundance of hydrophobic and polar residues in an individual sequence both within the full sequence and the LCR. We observed the correlation between the polar and hydrophobic residues to be less strongly pronounced within the LLPS-prone dataset compared to Swiss-Prot both on the level of the entire sequence (Figure 2c) and within the LCRs (Figure 2d), potentially due to the regions of polar residues being punctuated by only a fixed number of hydrophobic residues regardless of the length of the polar region. This observation was in line with the stickers-and-spacers model that has been proposed previously, whereby extended regions of polar amino acids patterned with hydrophobic residues govern the onset of protein LLPS.^22^

### Model for classifying the propensity of unseen sequences to phase separate

We next set out to develop machine learning classifiers for predicting the propensity of protein and peptide constructs to undergo LLPS using the constructed datasets for training. To featurise the protein sequences we used the aforedescribed physicochemical and amino acid sequence-specific features in combination with the word2vec sequence featurisation algorithm.^36^ Specifically, the word2vec embedding vectors were trained in an unsupervised manner on the full Swiss-Prot database using skip-gram pre-training procedure with 3-grams, a window size of 25 amino acids and negative sampling (see Methods).

We first trained two classifiers, Model A1 and Model A2, distinguishing sequences that phase separated at a low concentration (LLPS^+^) from fully ordered sequences highly unlikely to undergo LLPS (PDB*). To this effect, a random subset of 137 entries was sampled from the PDB* dataset ensuring that during the training process a comparable number of points from both datasets was encountered. Model A1 used 8 distinct physical features — the sequence length, hydrophobicity, Shannon entropy, the fraction of the sequence identified to be part of the LCRs and IDRs (Figure 1) and the fraction of polar, aromatic and cationic amino acid residues within the LCRs (Figure 2) for sequence featurisation. Model A2 featurised the sequences through 200-dimensional pre-trained word2vec embedding vectors. The feature vectors of all the data points were visualised on a 2D plane using t-distributed stochastic neighbour embedding for the physicochemical features (Figure 3a) and the word2vec-based (Figure 3b) embeddings. As desired, this dimensionality reduction process revealed a notable degree of separation into distinct clusters (LLPS^+^ – cyan, PDB* – magenta) for both featurisation approaches.

**Figure 3.**
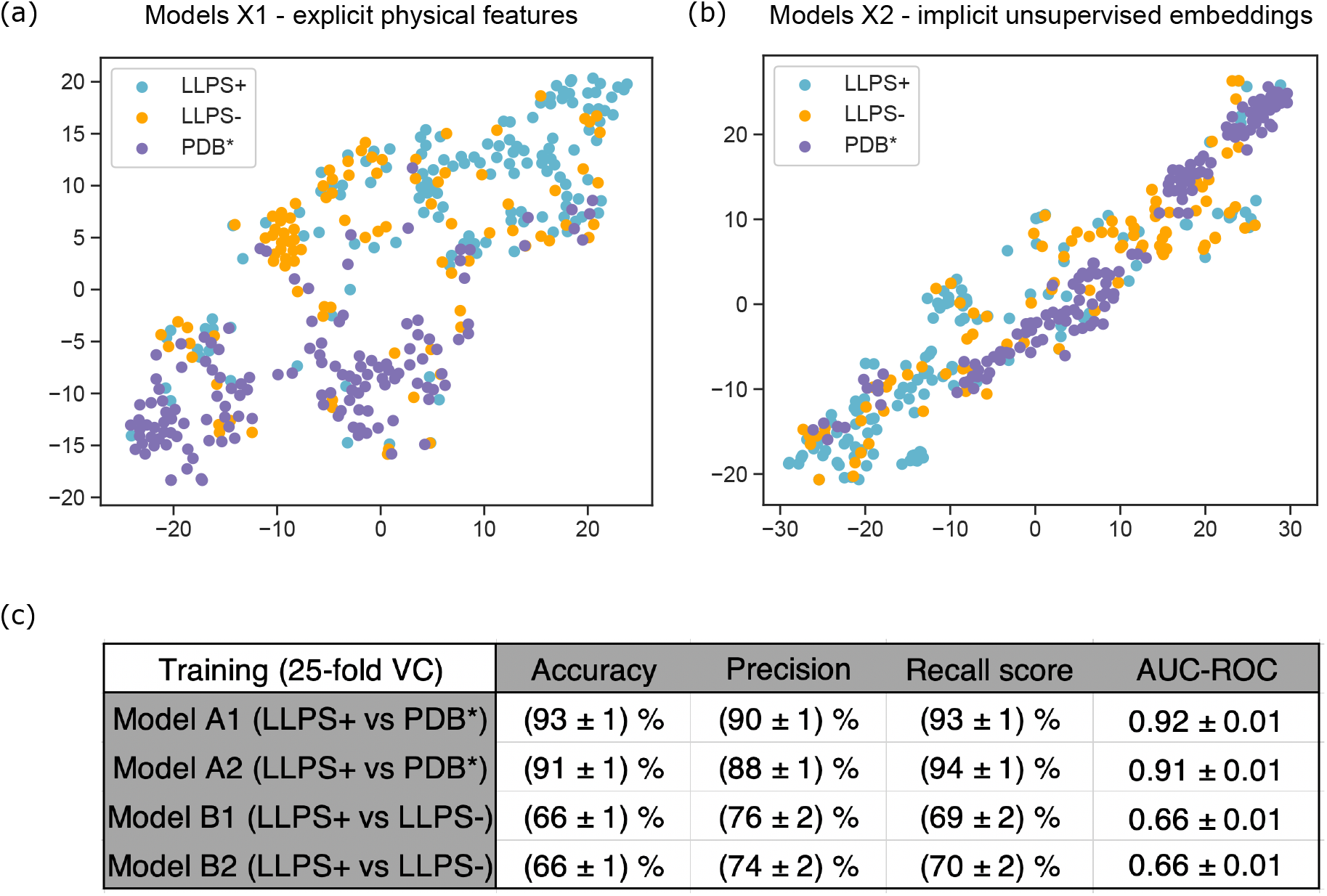
Cross-validation performance of the developed models. /-distributed stochastic neighbour embedding visualising the similarity of the **(a)** 8-dimensional vectors describing the physicochemical and amino acid composition specific features of the sequences and **(b)** 200-dimensional word2vec algorithm based embedding vectors for the LLPS^+^ (cyan), the LLPS^-^ (orange) and the PDB* (magenta) datasets on a 2D plane. **(c)** The performance of the four models on the training data with 25-fold cross-validation. The higher performance of Models A1 and A2 can be rationalised by the training sets (LLPS^+^ and PDB*) being more disjoint (Figure 1c-g) and the embedding vectors showing stronger level of clustering than is the case for the data used for training Models B1 and B2 (LLPS^+^ and LLPS^-^), both when physicochemical characteristics (panel **(a)**, Models A1 and B1) and implicit embedding vectors (panel **(b)**, Models A2 and B2) are used.

These data were used to train a model for classifying the data points into the two categories. To ensure generalisability, during the training process, the data were split into training and validation sets in a stratified manner, so that sequences with the same UniProt ID would belong to the same set. Using this strategy, a random forest classifier was trained (see Methods), which achieved over 93% accuracy and precision in correctly predicting the origin of sequences *(cv =* 25-fold cross-validation; Figure 3c), a result that is similar to a recently developed model.^31^

As the datasets on which Models A1 and A2 were trained were highly disjoint on the level of disorder of the individual sequences (Figure 1g), we proceeded by building additional models that could discriminate between the much more similar datasets LLPS^+^ and LLPS^-^ (Figure 1a). Such models could encompass information beyond disorder and have an enhanced potential to evaluate the propensity of different disordered proteins to undergo LLPS. As before, both the previously described physicochemical features (Model B1) and word2vec embeddings (Model B2) were used. However, this time, the resulting t-distributed stochastic neighbour embeddings were found to not cluster the points as effectively into the two distinct sets (Figure 3a–b). Indeed, a random forest classifier was found to achieve around 66% accuracy and 74–76% precision in correctly identifying the origin of the proteins in the validation data set (cv = 25-fold cross-validation; Figure 3c).

### Performance of the models on an external dataset

Having established the high cross-validation performance of the models within our generated datasets, we set out to test the model on external test data. To this effect, we used the PhaSepDB^32^ database, which had been curated by extracting all publications from the NCBI PubMed database that included phase separation related keywords in their abstracts. The resulting 2763 papers were manually rechecked to obtain publications that described membraneless organelles and related proteins, and they were further manually filtered to these proteins that had been observed to undergo phase separation experimentally either *in vitro* or *in vivo*. After removing from this database the UniProt IDs that overlapped with any of the LLPSDB entries and hence with our training data, we obtained a set of 289 LLPS-prone human sequences (Supplementary Table S4). A further examination of these 289 sequences highlighted that 40 of them included no intrinsically disorder regions. To develop an insight into the likelihood of fully ordered sequences undergoing phase separation, we analysed the LLPSDB database (that was used to construct the training datasets) listing all protein constructs together with the environmental conditions under which they had been studied. This analysis revealed that while some proteins with no disordered regions had been observed to phase separate, these experiments were performed under an extensive amount of molecular crowding (e.g. dextran, ficoll, polyethylene glycol) or in a non-homotypical environment (e.g. in the presence of lipids). It is thus likely that the fully-ordered sequences within the PhaSepDB that were identified as phase separating in a keyword-based literature search, similarly phase separated under non-homotypical conditions or notable level of molecular crowding, and their phase-transition cannot be directly linked to the protein sequence as it was not triggered exclusively by homotypic interactions between protein molecules. Motivated by this argument, we eliminated fully structured sequences, which yielded a list of 249 sequences serving as our external test data (Supplementary Table S5).

The predictions made by the four models on the human proteins of the Swiss-Prot dataset (20,291 entries; sequences that overlapped with the training set were excluded) are highlighted in Figures 4a-b with the sequences that are part of the external test data indicated as coloured points on the probability density function. By setting the classification threshold to 0.5, we observed that Models A1 and A2 (trained on LLPS^+^ and PDB*) identified 97% and 95% (Figure 4a, inset) and Models B1 and B2 (trained on LLPS^+^ and LLPS^-^) 81% and 83% (Figure 4b, inset) of the proteins in the external test data set correctly as phase-separating. We proceeded by developing an insight into the specificity-sensitivity relationships of the trained models. As the phase behaviour of many proteins remains unstudied, it is challenging to quantify the true false positive rate of any model predicting phase separating sequences. However, relying on the exhaustive keyword-based literature search performed as part of the construction of the PhaSepDB database and using the proteins that were not identified as LLPS-positive through this PubMed search we were able to define its upper bound. Using this approximation, we constructed the receiver operator characteristic (ROC) curves for the four models (Figures 4d-e), where the ordinate showed the fraction of the proteins in the test set that were identified correctly as LLPS-prone (true positive rate) and the abscissa showed the fraction of the remaining proteins in the human proteome predicted to be LLPS-positive at this threshold (upper bound for the false positive rate). These relationships indicated that while Models A1–2 exhibited a higher sensitivity and a lower false negative rate in identifying phase separating sequences compared to Models B1–2, they could also demonstrate a higher false positive rate and a lower specificity. This result was likely a direct effect of the training datasets used dor constructing Models A1–2 being highly disjoint on multiple levels (Figures 1–2), most notably on their degree of intrinsic disorder, which limited the possibility of the models to obtain insight into other sequence-determinants that define protein phase behaviour.

**Figure 4.**
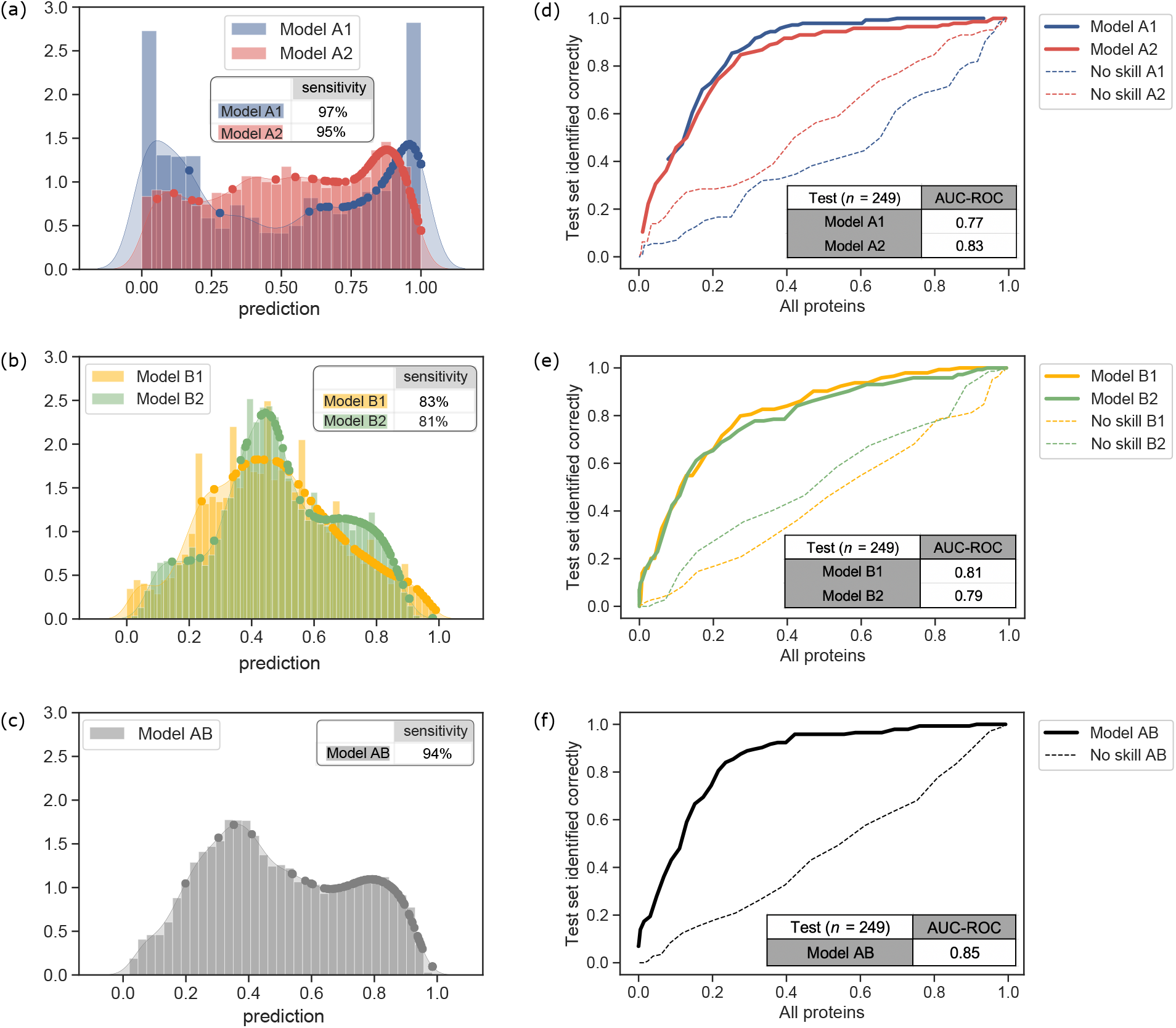
Performance of the models on external test data, including *n =* 249 sequences with disordered regions (Supplementary Table S5) from the PhaSepDB32 curated based on searching the NCBI PubMed database for LLPS-associated keywords. The prediction made by **(a)** Models A1–2, **(b)** Models B1–2 and **(c)** Model AB (ensemble classifier constructed as a mean of the four models) on the human sequences in the Swiss-Prot database (20,291 sequences) with the test proteins indicated as coloured points on the probability density function. All the models achieved a high sensitivity in correctly identifying the LLPS-prone proteins from the external data. Approximated receiver operator characteristic (ROC) curves for **(d)** Models A1–2, **(e)** Models B1–2 and **(f)** Model AB. Model AB achieved both high specificity and sensitivity in identifying LLPS-prone proteins. The “no skill” models (dotted lines) correspond to models generated by randomly shuffling the predictions between proteins.

Finally, with the developed models (A1–2, B1–2) exhibiting different desired features, we generated a final model (Model AB) where the prediction on every sequences was set to be the average prediction made on it by the four individual models. This combined model showed a high 94% sensitivity on identifying LLPS-prone seuqneces from the external test data. Crucially, the area under its approximated ROC-curve exceeded what could be achieved by either of the two strategies (training on LLPS^+^ vs PDB* or on LLPS^+^ vs LLPS^-^) on their own (Figure 4f). This optimal performance highlighted how a combination of individual weak predictors (here simply by estimating their mean, although more complex ensemble methods could be considered) has the potential to result in classifiers that can retain the best features of their constituent models.^37,38^ Sequences from this list (Supplementary Table S6) to which this combined model assigned a high LLPS propensity could be interesting candidates for experimental investigation of their phase behaviour.

### Conclusion

To understand how protein sequence governs its phase behaviour and further build an algorithm for predicting LLPS-prone sequences, we constructed datasets of proteins and peptides of varying LLPS propensity. The analysis of the curated datasets highlighted that LLPS-prone sequences were less hydrophobic, exhibited reduced entropy and had a higher degree of disorder than an average protein in the Swiss-Prot database. Furthermore, the data suggested that LLPS-prone proteins had long LCRs rich in polar amino acid residues and that the abundance of the latter residues correlated poorly with that of the hydrophobic residues within these regions, likely because the polar regions were punctuated by only a fixed number of hydrophobic residues regardless of the extent of the polar regions.^22^

Using the generated datasets and the identified characteristic sequence-specific features in combination with neural network based word2vec embeddings, we trained machine learning classifiers for predicting protein phase behaviour. Our models trained to distinguish LLPS-prone sequences from fully structured sequences (Models A1-2) achieved over 93% cross-validation accuracy in training and demonstrated over 97% sensitivity in identifying phase separating sequences from an external database of 249 LLPS-prone sequences. Combining their predictions with alternative models trained on sequences of varying LLPS propensity (Models B1-2) with the aim develop a more nuanced insight into the molecular features that govern protein phase behaviour, we built an improved classifier (Model AB) that showed both high specificity and sensitivity in identifying LLPS-positive proteins (approximated ROC–AUC of 0.85). Our results establish a model rooted in molecular principles for the description of protein phase behaviour from its sequence and highlight that the combination of individual weak predictors has the potential to create classifiers that retain best aspects from individual models.

## Methods

### Construction of the LLPS^+^ and LLPS^-^ datasets

The LLPS^+^ and LLPS^-^ datasets were constructed using the previously published LLPSDB database (accessed on 20 May 2020).^28^ Specifically, the “LLPS_Natural_protein” repository from “Datasets classified by protein name” was used, which documented a total of 2143 entries of proteins and their constructs examined for the occurrence of LLPS under various experimental conditions. The 2143 entries were filtered down to systems that included only a single naturally occurring protein with no post-translational modifications, repeat or single site mutations and to experiments where the examined protein sequence was longer than 50 amino acids. This procedure resulted in a dataset with 769 experimental entries including a total of 231 unique constructs from 120 different UniProt IDs.

For each of the constructs, the experiments where the construct had been observed to phase separate were combined and the average concentration at which these positive experiments were performed was evaluated. When the latter concentration was below 100 μM the construct was regarded LLPS-prone and it was included in the LLPS^+^ dataset (Supplementary Table S1). This process resulted in a total of 137 of such constructs from 77 unique Uniprot IDs. The constructs that had been observed to phase separate at a concentration higher than 100 μM were combined with the constructs that had not been observed to phase separate to generate the LLPS^-^ dataset. The latter dataset included a total of 94 entries from 61 unique UniProt IDs.

### Construction of the PDB^*^ dataset

Entries in the Protein Data Bank (PDB)^29^ were used to generate a diverse set of proteins highly unlikely to undergo LLPS. Specifically, first, amino acid chains that were fully structured (i.e., did not include any disordered residues) were extracted, which resulted in a total of 112,572 chains. PDB chains were matched to their corresponding UniProt IDs using SIFTS (EBI), and entries where sequence length did not match were discarded. Duplicate entries were removed and the remaining 13,325 chains were clustered for their sequence identity using a conservative cut-off of 30%. One sequence from each cluster was selected, resulting in the final dataset of 1563 sequences.

### Estimation of physical features from the sequences

A range of explicit physicochemical features was extracted for all the sequences in the four datasets from their amino acid sequences (LLPS^+^, LLPS^-^, PDB* and Swiss-Prot). Specifically, the molecular weight of each sequence and its amino acid composition were calculated using Python package BioPython. The hydrophobicity of each of the sequence was evaluated by summing the individual hydrophobicity values of the amino acids in the sequences using the Kyte and Doolittle hydropathy scale.^33^ The Shannon entropy of each sequence was estimated from the following formula:

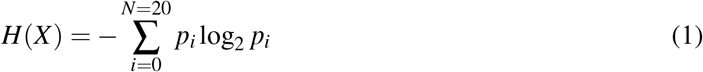

where *p* corresponds to the frequency of each of the naturally occurring twenty amino acid in the sequence. The low complexity regions (LCR) for each of the sequences was estimated using the SEG algorithm^34^ with standard parameters. The disordered region was predicted with IUPred2a^35^ that estimated the probability of disorder for each of the individual amino acid residues in the sequence. The disorder fraction of a sequence was calculated as the fraction of residues in the total sequence that were considered disordered where a specific residue was classified as disordered when the disorder probability stayed above 0.5 for at least 20 consecutive residues.

Finally, the amino acid sequence and the LCR regions were described for their amino acid content by allocating the residues to the following groups: amino acids with polar residues (Serine, Glutamine, Asparagine, Glycine, Cysteine, Threonine, Proline), with hydrophobic residues (Alanine, Isoleucine, Leucine, Methionine, Phenylalanine, Valine), with aromatic residues (Truptophan, Tyrosine, Phenylalanine), with cationic residues (Lysine, Arginine, Histidine) and with anionic residues (Aspartic acid, Glutamic acid).

### Protein sequence embeddings

Unsupervised protein sequence embeddings were used for featurising all the analysed protein sequences. Specifically, word2vec skip-gram pre-training procedure was used with the help of Python gensim library^39^ to generate 200-dimensional embedding vectors. The pre-training was performed on the full Swiss-Prot database (accessed on 26 Jun 2020) using 3-grams, a window size of 25 and negative sampling. This process resulted in 200-dimensional embedding vectors for each of the sequences that served as the input features when training the machine learning classifiers. Each protein sequence was broken into 3-grams using all three possible reading frames. The final 200-dimensional full protein embedding was obtained by summing all constituent 3-gram embeddings.

### Machine learning classifier training and performance estimation

Two classifiers were built based on the trained implicit embedding vectors. Specifically, Models A1–2 were trained to differentiate the sequences in the LLPS^+^ dataset from those in the PDB* and Models B1–2 were trained to differentiate between the sequences in the LLPS^+^ and LLPS^+^ datasets. For the first case, 137 entries from the total of 1563 sequences in the PDB^*^ were sampled at random to generate a balanced dataset between the classes. For performing the training, the dataset was split into train and validation test in 1: 4 ratio and 25-fold was cross-validation used to estimate the performance of the model. For each fold, the split was performed in a stratified manner, such that entries from the same UniProt ID were always grouped together either to the training set or the validation set. A random seed of 42 was used throughout.

For all the four models, a random forest classifier was constructed using Python scikit-learn package^40^ with default parameters. No hyperparameter tuning was performed for either of the cases — while such a tuning step may have given an improvement in accuracy it can also lead to an overfitted model that does not generalise well to unseen data. All “no skill” models were constructed by randomly shuffling the predictions made by this model on all the proteins of interest.

## Acknowledgements

The research leading to these results has received funding from the Schmidt Science Fellows program in partnership with the Rhodes Trust (K.L.S.), St John’s College Junior Research Fellowship (K.L.S.), Trinity College Krishnan-Ang Studentship (R.Q.) and the Honorary Trinity-Henry Barlow Scholarship (R.Q.), the EPSRC Cambridge NanoDTC (EP/L015978/1; W.E.A.), the EPSRC Impact Acceleration Programme (W.E.A., G.K., T.P.J.K.), the European Research Council under the European Union’s Horizon 2020 Framework Programme through the Marie Sklodowska-Curie grant MicroSPARK (agreement n° 841466; G.K.), the Herchel Smith Fund of the University of Cambridge (G.K.), Wolfson College Junior Research Fellowship (G.K.), the European Research Council under the European Union’s Seventh Framework Programme (FP7/2007–2013) through the ERC grants PhysProt (agreement n^°^ 337969; T.P.J.K) and the Newman Foundation (T.P.J.K.).

## Author contributions

K.L.S., A.S.M. and T.P.J.K. designed and conceptualised the study, K.L.S and A.S.M. performed the analysis and R.Q. built the web interface. K.L.S., A.S.M. and T.P.J.K. wrote the manuscript, all authors reviewed the manuscript and commented on it.

## References

[1] C. P. Brangwynne, C. R. Eckmann, D. S. Courson, A. Rybarska, C. Hoege, J. Gharakhani, F. Jülicher, and A. A. Hyman, “Germline P granules are liquid droplets that localize by controlled dissolution/condensation,” Science, vol. 324, no. 5935, pp. 1729–1732, 2009.

[2] A. A. Hyman and C. P. Brangwynne, “Beyond stereospecificity: liquids and mesoscale organization of cytoplasm,” Developmental Cell, vol. 21, no. 1, pp. 14–16, 2011.

[3] A. A. Hyman, C. A. Weber, and F. Jülicher, “Liquid–liquid phase separation in biology,” Annual Review of Cell and Developmental Biology, vol. 30, pp. 39–58, 2014.

[4] S. Boeynaems, S. Alberti, N. L. Fawzi, T. Mittag, M. Polymenidou, F. Rousseau, J. Schymkowitz, J. Shorter, B. Wolozin, L. Van Den Bosch, et al., “Protein phase separation: a new phase in cell biology,” Trends in Cell Biology, vol. 28, no. 6, pp. 420–435, 2018.

[5] S. Alberti and D. Dormann, “Liquid–liquid phase separation in disease,” Annual Review of Genetics, vol. 53, pp. 171–194, 2019.

[6] A. Aguzzi and M. Altmeyer, “Phase separation: linking cellular compartmentalization to disease,” Trends in Cell Biology, vol. 26, no. 7, pp. 547–558, 2016.

[7] P. Li, S. Banjade, H.-C. Cheng, S. Kim, B. Chen, L. Guo, M. Llaguno, J. V. Hollingsworth, D. S. King, S. F. Banani, et al., “Phase transitions in the assembly of multivalent signalling proteins,” Nature, vol. 483, no. 7389, pp. 336–340, 2012.

[8] E. Sokolova, E. Spruijt, M. M. Hansen, E. Dubuc, J. Groen, V. Chokkalingam, A. Piruska, H. A. Heus, and W. T. Huck, “Enhanced transcription rates in membrane-free protocells formed by coacervation of cell lysate,” Proceedings of the National Academy of Sciences, vol. 110, no. 29, pp. 11692–11697, 2013.

[9] J. Berry, S. C. Weber, N. Vaidya, M. Haataja, and C. P. Brangwynne, “RNA transcription modulates phase transition-driven nuclear body assembly,” Proceedings of the National Academy of Sciences, vol. 112, no. 38, pp. E5237–E5245, 2015.

[10] Y. Shin and C. P. Brangwynne, “Liquid phase condensation in cell physiology and disease,” Science, vol. 357, no. 6357, 2017.

[11] J. A. Riback, C. D. Katanski, J. L. Kear-Scott, E. V. Pilipenko, A. E. Rojek, T. R. Sosnick, and D. A. Drummond, “Stress-triggered phase separation is an adaptive, evolutionarily tuned response,” Cell, vol. 168, no. 6, pp. 1028–1040, 2017.

[12] C. P. Brangwynne, P. Tompa, and R. V. Pappu, “Polymer physics of intracellular phase transitions,” Nature Physics, vol. 11, no. 11, pp. 899–904, 2015.

[13] E. Gomes and J. Shorter, “The molecular language of membraneless organelles,” Journal of Biological Chemistry, vol. 294, no. 18, pp. 7115–7127, 2019.

[14] A. S. Raut and D. S. Kalonia, “Effect of excipients on Liquid–liquid phase separation and aggregation in dual variable domain immunoglobulin protein solutions,” Molecular Pharmaceutics, vol. 13, no. 3, pp. 774–783, 2016.

[15] K. Julius, J. Weine, M. Gao, J. Latarius, M. Elbers, M. Paulus, M. Tolan, and R. Winter, “Impact of macromolecular crowding and compression on protein-protein interactions and Liquid–liquid phase separation phenomena,” Macromolecules, vol. 52, no. 4, pp. 1772–1784, 2019.

[16] L. Lemetti, S.-P. Hirvonen, D. Fedorov, P. Batys, M. Sammalkorpi, H. Tenhu, M. B. Linder, and A. S. Aranko, “Molecular crowding facilitates assembly of spidroin-like proteins through phase separation,” European Polymer Journal, vol. 112, pp. 539–546, 2019.

[17] G. Krainer, T. J. Welsh, J. A. Joseph, J. R. Espinosa, E. de Csillery, A. Sridhar, Z. Toprakcioglu, G. Gudiskyte, M. A. Czekalska, W. E. Arter, et al., “Reentrant liquid condensate phase of proteins is stabilized by hydrophobic and non-ionic interactions,” bioRxiv, 2020.

[18] E. W. Martin and T. Mittag, “Relationship of sequence and phase separation in protein low-complexity regions,” Biochemistry, vol. 57, no. 17, pp. 2478–2487, 2018.

[19] S. Alberti, A. Gladfelter, and T. Mittag, “Considerations and challenges in studying Liquid–liquid phase separation and biomolecular condensates,” Cell, vol. 176, no. 3, pp. 419–434, 2019.

[20] R. M. Vernon, P. A. Chong, B. Tsang, T. H. Kim, A. Bah, P. Farber, H. Lin, and J. D. Forman-Kay, “Pi-pi contacts are an overlooked protein feature relevant to phase separation,” Elife, vol. 7, p. e31486, 2018.

[21] G. L. Dignon, R. B. Best, and J. Mittal, “Biomolecular phase separation: From molecular driving forces to macroscopic properties,” Annual Review of Physical Chemistry, vol. 71, pp. 53–75, 2020.

[22] E. W. Martin, A. S. Holehouse, I. Peran, M. Farag, J. J. Incicco, A. Bremer, C. R. Grace, A. Soranno, R. V. Pappu, and T. Mittag, “Valence and patterning of aromatic residues determine the phase behavior of prion-like domains,” Science, vol. 367, no. 6478, pp. 694–699, 2020.

[23] J. Wang, J.-M. Choi, A. S. Holehouse, H. O. Lee, X. Zhang, M. Jahnel, S. Maharana, R. Lemaitre, A. Pozniakovsky, D. Drechsel, et al., “A molecular grammar governing the driving forces for phase separation of prion-like RNA binding proteins,” Cell, vol. 174, no. 3, pp. 688–699, 2018.

[24] R. M. Vernon and J. D. Forman-Kay, “First-generation predictors of biological protein phase separation,” Current Opinion in Structural Biology, vol. 58, pp. 88–96, 2019.

[25] H.-R. Li, W.-C. Chiang, P.-C. Chou, W.-J. Wang, and J.-r. Huang, “Tar dna-binding protein 43 (tdp-43) Liquid–liquid phase separation is mediated by just a few aromatic residues,” Journal of Biological Chemistry, vol. 293, no. 16, pp. 6090–6098, 2018.

[26] S. Elbaum-Garfinkle, Y. Kim, K. Szczepaniak, C. C.-H. Chen, C. R. Eckmann, S. Myong, and C. P. Brangwynne, “The disordered P granule protein LAF-1 drives phase separation into droplets with tunable viscosity and dynamics,” Proceedings of the National Academy of Sciences, vol. 112, no. 23, pp. 7189–7194, 2015.

[27] T. P. Dao, R.-M. Kolaitis, H. J. Kim, K. O’Donovan, B. Martyniak, E. Colicino, H. Hehnly, J. P. Taylor, and C. A. Castañeda, “Ubiquitin modulates Liquid–liquid phase separation of UBQLN2 via disruption of multivalent interactions,” Molecular Cell, vol. 69, no. 6, pp. 965–978, 2018.

[28] Q. Li, X. Peng, Y. Li, W. Tang, J. Zhu, J. Huang, Y. Qi, and Z. Zhang, “LLPSDB: a database of proteins undergoing Liquid–liquid phase separation in vitro,” Nucleic Acids Research, vol. 48, no. D1, pp. D320–D327, 2020.

[29] H. M. Berman, J. Westbrook, Z. Feng, G. Gilliland, T. N. Bhat, H. Weissig, I. N. Shindyalov, and P. E. Bourne, “The protein data bank (accessed on 26 jun 2020),” Nucleic Acids Research, vol. 28, no. 1, pp. 235–242, 2000.

[30] UniProt Consortium, “UniProt: a worldwide hub of protein knowledge,” Nucleic Acids Research, vol. 47, no. D1, pp. D506–D515, 2019.

[31] T. Sun, Q. Li, Y. Xu, Z. Zhang, L. Lai, and J. Pei, “Prediction of Liquid–liquid phase separation proteins using machine learning,” bioRxiv, 2019.

[32] K. You, Q. Huang, C. Yu, B. Shen, C. Sevilla, M. Shi, H. Hermjakob, Y. Chen, and T. Li, “PhaSepDB: a database of Liquid–liquid phase separation related proteins,” Nucleic Acids Research, vol. 48, no. D1, pp. D354–D359, 2020.

[33] J. Kyte and R. F. Doolittle, “A simple method for displaying the hydropathic character of a protein,” Journal of Molecular Biology, vol. 157, no. 1, pp. 105–132, 1982.

[34] J. C. Wootton and S. Federhen, “Statistics of local complexity in amino acid sequences and sequence databases,” Computers & Chemistry, vol. 17, no. 2, pp. 149–163, 1993.

[35] Z. Dosztanyi, V. Csizmok, P. Tompa, and I. Simon, “The pairwise energy content estimated from amino acid composition discriminates between folded and intrinsically unstructured proteins,” Journal of Molecular Biology, vol. 347, no. 4, pp. 827–839, 2005.

[36] T. Mikolov, K. Chen, G. Corrado, and J. Dean, “Efficient estimation of word representations in vector space,” arXiv preprint arXiv:1301.3781, 2013.

[37] L. Rokach, “Ensemble-based classifiers,” Artificial Intelligence Review, vol. 33, no. 1-2, pp. 1–39, 2010.

[38] S. B. Kotsiantis, I. D. Zaharakis, and P. E. Pintelas, “Machine learning: a review of classification and combining techniques,” Artificial Intelligence Review, vol. 26, no. 3, pp. 159–190, 2006.

[39] R. Řehuůek and P. Sojka, “Software Framework for Topic Modelling with Large Corpora,” in Proceedings of the LREC 2010 Workshop on New Challenges for NLP Frameworks, (Valletta, Malta), pp. 45–50, ELRA, May 2010. http://is.muni.cz/publication/884893/en.

[40] F. Pedregosa, G. Varoquaux, A. Gramfort, V. Michel, B. Thirion, O. Grisel, M. Blondel, P. Prettenhofer, R. Weiss, V. Dubourg, et al., “Scikit-learn: Machine learning in python,” Journal of Machine Learning Research, vol. 12, no. Oct, pp. 2825–2830, 2011.

